## Supplementary Information for "Sampling-based Bayesian inference in recurrent circuits of stochastic spiking neurons"

### 1 Suplementary Figures

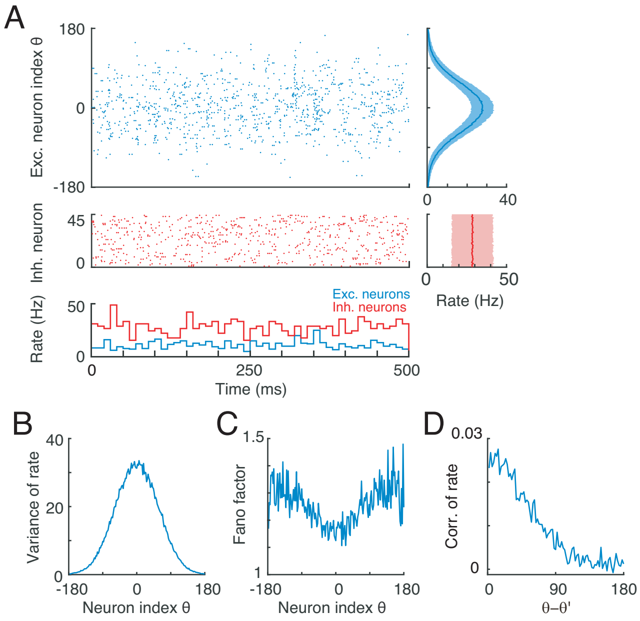

Figure S1: (A) Example realization of population spiking activity of excitatory and inhibitory neurons in a single recurrent network as shown in Figure 1 in the main text. Right: the firing rate of neurons averaged over time, with shaded region encompassing one standard deviation from the mean. (B-C) The variance of firing rate (B) and Fano factor (C) of responses of E neurons in the network. (D) The spike count correlation between two excitatory (E) neurons with their difference of preferred stimulus features. Parameters: $U^{\mathrm{f}} = 30$Hz, $w_E = 4 \times 10^{-3}$, and others are the same as the ones listed in Table 1.

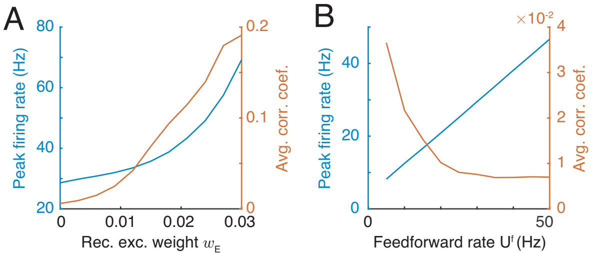

Figure S2: In the single recurrent network shown in Figure 4, the peak firing rate (blue) and the averaged correlation coefficient (red) of E neurons in the network model as a function of recurrent excitatory weight (A) and the feedforward input rate (B). Parameters: (A) $U^{\mathrm{f}} = 30$Hz, (B) $w_E = 2 \times 10^{-3}$, and others are the same as the one listed in Table 1.

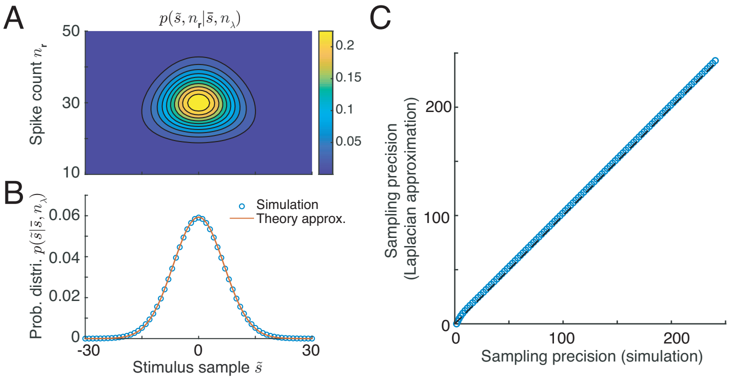

Figure S3: The sampling distribution of stimulus from internal Poisson spike generation given a Gaussian profile population firing rate. (A) The joint distribution of stimulus samples and spike counts of all neurons given a Gaussian profile population firing rate (Eq. S3). (B) The sampling distribution averaged from different values of spike counts. (C) The sampling precision (inverse of sampling variability) obtained using a Laplacian approximation (y-axis) and numerical simulation (x-axis).

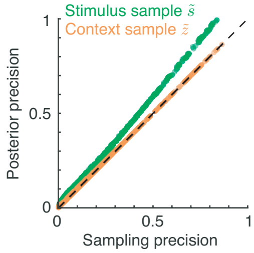

Figure S4: The comparison of the sampling precision of stimulus $s$ and context $z$ in a single recurrent network (shown in Figure 4) with posterior precision under different combinations of feedforward input rate and recurrent weight. Due to the negative rectification of the multiplicative variability in the recurrent inputs (Eq. 48), the precision of stimulus samples is lower than the posterior precision especially at large recurrent input strength (the dots with high precision) where the rectification effect is strong. In the simulation, the feedforward input rate $U^{\mathsf{f}}$ and recurrent weight $w_E$ were randomly generated. Each dot in the figure is the result under a particular combination of randomly generated $U^{\mathsf{f}}$ and $w_E$, and there are 1000 dots in above figure. Parameters: $U^{\mathsf{f}} \in \mathcal{U}(0, 50)$Hz, $w_E \in \mathcal{U}(0.001, 0.051)$ where $\mathcal{U}(a, b)$ represents a uniform distribution ranges from $a$ to $b$. The other parameters of the network are the same as the ones listed in Table 1.

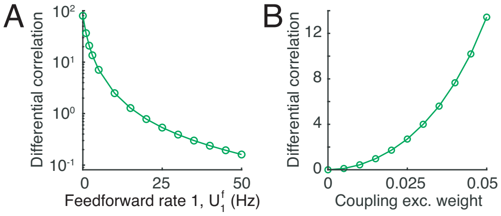

Figure S5: In the coupled networks shown in Figure 6, the amount of internally generated differential correlations due to internal spike generation as a function of feedforward rate applied to network 1 (A), and the coupling weight between two networks (B). (A) Increasing the firing rate of feedforward input 1 decreases the amount of internally generated differential correlations in network 1. (B) Increasing the coupling weight between two networks increases the internally generated differential correlation. Parameters: (A) $w_{EE}^{12} = w_{EE}^{21} = 0.01$, (B) $U^{\mathsf{f}} = 30$Hz.

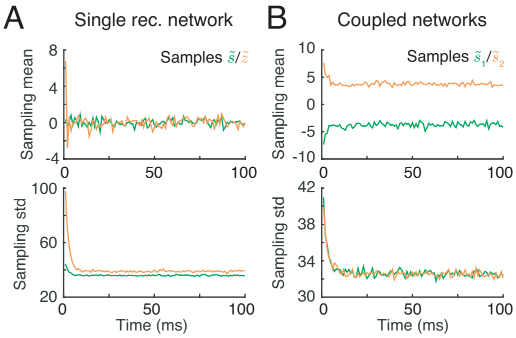

Figure S6: The sampling paths averaged from 10,000 different realizations in a single recurrent network as shown in Figure 1 (A), and in coupled networks as shown in Figure 6 (B). The sampling-based inference in stimulus subspace converges in less than 20ms. Parameters: (A) $w_E = 4 \times 10^{-3}$, $U^{\mathrm{f}} = 30$Hz; (B) $w_{EE}^{12} = w_{EE}^{21} = 0.01$, $U^{\mathrm{f}1} = U^{\mathrm{f}2} = 30$Hz, and others are the same as the ones listed in Table 1.

### 2 Internal Poisson spiking variability samples stimulus

#### 2.1 The sampling distribution under Gaussian profile firing rate

Here we show analytically that Poisson variability of internal spike generation provides the appropriate variability to drive sampling from a specified continuous distribution. By substituting the instantaneous firing rate $\boldsymbol{\lambda}_t$ (Eq. 12, which is copied in below for ease of exposition)

$$\boldsymbol{\lambda}_{tj} = R \exp[-(\bar{s}_t - \theta_j)^2/2a^2]$$

into the expression for Poisson spike generation over the network (Eq. 11),

$$\mathbf{r}_t \quad \sim \quad \textstyle\prod_{j=1}^{N_E} \text{Poisson}\left(\boldsymbol{\lambda}_{tj}\Delta t\right),$$

the probability of observed population spikes becomes (for concise notations we suppress the time index $t$ in the derivation below),

$$\begin{aligned} p\left(\mathbf{r}|\boldsymbol{\lambda}\right) &= \textstyle\prod_{j=1}^{N_E} \text{Poisson}\left(\mathbf{r}_j|\boldsymbol{\lambda}_j\Delta t\right), \\ &\propto (R\Delta t)^{\sum_j r_j} \exp\left[-\textstyle\sum_j r_j \frac{(\bar{s}-\theta_j)^2}{2a^2}\right] \exp\left[-R\Delta t \textstyle\sum_j e^{-(\bar{s}-\theta_j)^2/2a^2}\right]. \end{aligned} \tag{S1}$$

To simplify notations, let

$$n_{\mathbf{r}} = \textstyle\sum_j \mathbf{r}_j, \quad n_{\boldsymbol{\lambda}} = \textstyle\sum_j \boldsymbol{\lambda}_j \Delta t = R\Delta t \textstyle\sum_j e^{-(\bar{s}-\theta_j)^2/2a^2}, \tag{S2}$$

so that $n_{\mathbf{r}}$ is the number of emitted spikes of the whole neural population, and $n_{\boldsymbol{\lambda}}$ is the sum of instantaneous firing rate of all neurons. With the uniform distribution of the neurons' preferred stimulus, i.e., $\{\theta_j\}_{j=1}^N$ is uniformly distributed, $n_{\boldsymbol{\lambda}}$ will be a constant irrespective of the value of $\bar{s}$.

Normalizing the distribution over spike counts in Eq. (S1) into a likelihood over $\bar{s}$ and $n_{\boldsymbol{\lambda}}$ (Fig. S3A) gives,

$$\begin{aligned} p(\mathbf{r}|\boldsymbol{\lambda}) &\propto p(\tilde{s}, n_{\mathbf{r}}|\bar{s}, n_{\boldsymbol{\lambda}}), \\ &\propto \mathcal{N}\left(\tilde{s}|\bar{s}, a^2 n_{\mathbf{r}}^{-1}\right) \text{Poisson}(n_{\mathbf{r}}|n_{\boldsymbol{\lambda}}), \end{aligned} \tag{S3}$$

where

$$\tilde{s} = \textstyle\sum_j \mathbf{r}_j \theta_j / \sum_j \mathbf{r}_j, \quad \bar{s} = \sum_j \boldsymbol{\lambda}_j \theta_j / \sum_j \boldsymbol{\lambda}_j, \tag{S4}$$

as presented in Eq. 14. Here $\tilde{s}$ is regarded as a stimulus sample.

When $\boldsymbol{\lambda}$ repeatedly generates spikes over time, the sampling distribution of $\tilde{s}$ can be calculated by marginalizing the likelihood (Eq. 13, last line) over different values of $n_{\mathbf{r}}$.

$$\begin{aligned} p(\tilde{s}|\boldsymbol{\lambda}) &= \sum_{n_{\mathbf{r}}} p(\tilde{s}, n_{\mathbf{r}}|\bar{s}, n_{\boldsymbol{\lambda}}), \\ &= \sum_{n_{\mathbf{r}}} \mathcal{N}\left(\tilde{s}|\bar{s}, a^2 n_{\mathbf{r}}^{-1}\right) \text{Poisson}(n_{\mathbf{r}}|n_{\boldsymbol{\lambda}}), \end{aligned} \tag{S5}$$

We use Laplace's method to approximate the marginalization in order to get an analytical expression ([1], see Appendix 7.1). To simplify the calculation, we approximate the Poisson distribution in the above equation by a Gaussian distribution, which works well when the summed firing rate $n_{\boldsymbol{\lambda}}$, is large.

$$\begin{aligned} p(\tilde{s}|\boldsymbol{\lambda}) &\approx \int \mathcal{N}\left(\tilde{s}|\bar{s}, a^2 n_{\mathbf{r}}^{-1}\right) \mathcal{N}(n_{\mathbf{r}}|n_{\boldsymbol{\lambda}}, n_{\boldsymbol{\lambda}}) dn_{\mathbf{r}}, \\ &\approx \mathcal{N}\left(\tilde{s}|\bar{s}, a^2 \hat{n}_{\mathbf{r}}^{-1}\right) \mathcal{N}(\hat{n}_{\mathbf{r}}|n_{\boldsymbol{\lambda}}, n_{\boldsymbol{\lambda}}) \det(\mathrm{H}/2\pi)^{-1/2}, \end{aligned} \tag{S6}$$

where the second approximation in the above equation is obtained using Laplace's method, and

$$\begin{aligned} \hat{n}_{\mathbf{r}} &= \arg\max_{n_{\mathbf{r}}} p(\tilde{s}, n_{\mathbf{r}}|\bar{s}, n_{\boldsymbol{\lambda}}) = n_{\boldsymbol{\lambda}}, \\ \mathrm{H} &= -\frac{\partial^2}{\partial n_{\mathbf{r}}^2} \ln p(\tilde{s}, n_{\mathbf{r}}|\bar{s}, n_{\boldsymbol{\lambda}})\Big|_{n_{\mathbf{r}}=\hat{n}_{\mathbf{r}}}. \end{aligned} \tag{S7}$$

After some tedious analytical calculations (see Appendix 7.1), the sampling distribution obtained over time given a fixed firing rate $\boldsymbol{\lambda}_t$ is (Fig. S3B),

$$p(\tilde{s}|\boldsymbol{\lambda}) \approx \mathcal{N}\left(\tilde{s}|\bar{s}, a^2 n_{\boldsymbol{\lambda}}^{-1}\right), \tag{S8}$$

where the mean, $\bar{s}$, and variance, $a^2 n_{\boldsymbol{\lambda}}^{-1}$, of the sampling distribution are both linear projections of the firing rate $\boldsymbol{\lambda}$ (Eqs. S2 and S4). Therefore the firing rate fully determines the sampling distribution, and the sum of the firing rate determines the sampling variability.

Fig. S3 numerically verifies that the sampling distribution indeed can be approximated by a Gaussian distribution. In particular, the sampling variability computed by Laplacian approximation is consistent with our numerical simulations (Fig. S3C).

#### 2.2 The sampling distribution under an arbitrary profile of population firing rate

Our above derivations are based on a Gaussian profile for the firing rate (Eq. 12). Here we extend our theory to a more general case. Similar to the assumption made in Eq. (12), we suppose the instantaneous firing rate, $\boldsymbol{\lambda}$ (time index $t$ is suppressed), is parameterized as

$$\boldsymbol{\lambda}_j = R\exp[\mathbf{h}_j(\bar{s})], \tag{S9}$$

where $\mathbf{h}_j(\bar{s})$ determines the profile of population firing rate. The same firing rate $\boldsymbol{\lambda}$ will be used to generate independent Poisson spikes across time. Substituting Eq. (S9) into the expression of Poisson spike generation (Eq. 11), and taking the logarithm of the distribution yields,

$$\ln p\,(\mathbf{r}|\boldsymbol{\lambda}) = \textstyle\sum_j \mathbf{r}_j \mathbf{h}_j(\bar{s}) + (\sum_j \mathbf{r}_j)\ln(R\Delta t) - R\Delta t \sum_j \exp[\mathbf{h}_j(\bar{s})] - \ln(\mathbf{r}_j!). \tag{S10}$$

Assuming that the preferred stimuli of the neurons in the network uniformly cover the stimulus domain, then for large $N_E$ the sum of population firing rate $R\Delta t \sum_j \exp[\mathbf{h}_j(\bar{s})]$ (the third term on the right hand side in above equation) will be a constant irrespective of the stimulus. Since the second and fourth terms on the right hand side of Eq. (S10) do not depend on $\bar{s}$ then the likelihood of $\bar{s}$ given an emitted population response $\mathbf{r}$ can be simplified to

$$\begin{aligned} p(\bar{s}|\mathbf{r}) &\propto \exp\left[\textstyle\sum_j \mathbf{r}_j \mathbf{h}_j(\bar{s})\right], \\ &\propto \exp\left[\mathbf{h}(\bar{s})^\top \mathbf{r}\right]. \end{aligned} \tag{S11}$$

Eq. (S11) has the same form as a probabilistic population code (PPC) [2], where the likelihood over $\bar{s}$ belongs to an exponential family of distributions. In particular, the logarithm of the population firing profile, $\mathbf{h}(\bar{s})$ (Eq. S9), determines the type of the likelihood of $\bar{s}$.

Instead of using a single observation of neuronal response, $\mathbf{r}$, to parametrically encode the whole likelihood of $\bar{s}$ as in a PPC [2], here we treat each $\mathbf{r}$ as a stimulus sample $\tilde{s}$ which is randomly drawn from the likelihood. The two different views of the population response $\mathbf{r}$ correspond to interchanging the random variable and the conditional parameters in the distribution,

$$\tilde{s} \sim p(\tilde{s}|\mathbf{r}) \propto \exp\left[\mathbf{h}(\tilde{s})^\top \mathbf{r}\right]. \tag{S12}$$

The stimulus sample $\tilde{s} = D(\mathbf{r})$ can be read out from $\mathbf{r}$ by a decoder $D$, whose detailed form depends on the population firing rate profile, $\mathbf{h}(s)$.

When using a fixed population firing rate $\boldsymbol{\lambda}$ with a known $\bar{s}$ (Eq. S9) to repeatedly generate a vector of spike counts, $\mathbf{r}$, across time, the distribution of stimulus samples collected over time is

$$\mathbb{E}_{p(\mathbf{r}|\boldsymbol{\lambda})}\left[p(\tilde{s}|\mathbf{r})\right] = \int p(\tilde{s}|\mathbf{r})p(\mathbf{r}|\boldsymbol{\lambda})d\mathbf{r}. \tag{S13}$$

Since the mean of the neuronal response is the same as the underlying firing rate, i.e., $\langle\mathbf{r}\rangle = \boldsymbol{\lambda}$, we assume that the distribution in the integrand of the above equation, i.e., $p(s|\mathbf{r})p(\mathbf{r}|\boldsymbol{\lambda})$, has a peak at the point where $\mathbf{r} = \boldsymbol{\lambda}$. With this assumption and using the Laplacian approximation, the distribution of stimulus samples collected over time becomes,

$$\begin{aligned}\mathbb{E}_{p(\mathbf{r}|\boldsymbol{\lambda})}\left[p(\tilde{s}|\mathbf{r})\right] &\propto \mathbb{E}_{p(\mathbf{r}|\boldsymbol{\lambda})}\left\{\exp\left[\mathbf{h}(\tilde{s})^\top\mathbf{r}\right]\right\}, \\ &\approx \exp\left[\mathbf{h}(\tilde{s})^\top\boldsymbol{\lambda}\right] \equiv p(\tilde{s}|\boldsymbol{\lambda}). \end{aligned}\tag{S14}$$

This implies that each stimulus sample is randomly drawn from a conditional distribution which is determined by the population firing rate profile,

$$\tilde{s} \sim p(\tilde{s}|\boldsymbol{\lambda}) \propto \exp\left[\mathbf{h}(\tilde{s})^\top\boldsymbol{\lambda}\right], \tag{S15}$$

which is shown in Eq. (16) of the main text.

### 3 The equilibrium of Gibbs sampling dynamics

#### 3.1 Hierarchically generative model

The iteration of the Gibbs sampling algorithm can be better analyzed by converting it into a discrete dynamical system. We rewrite the Gibbs sampling dynamics (Eqs. 25-26 in maintext) here to ease the exposition:

$$\begin{aligned} p(s|\tilde{z}_t, \mathbf{u}^\mathsf{f}) &\propto \mathcal{N}\left(s|\bar{s}_t, \Lambda^{-1}\right), & \text{(S16)}\\ \bar{s}_t &= \frac{\Lambda_\mathsf{f}\mu_\mathsf{f} + \Lambda_s\tilde{z}_t}{\Lambda_\mathsf{f}+\Lambda_s}, \quad \Lambda = \Lambda_\mathsf{f} + \Lambda_s. \\ \tilde{s}_t &\sim \mathcal{N}\left(s|\bar{s}_t, \Lambda^{-1}\right). & \text{(S17)}\\ \tilde{z}_{t+\Delta t} &\sim \mathcal{N}(z|\tilde{s}_t, \Lambda_s^{-1}). & \text{(S18)}\end{aligned}$$

The above three steps in the Gibbs sampling algorithm can be written as the iterations of the following discrete dynamical system,

$$\begin{aligned}\bar{s}_t &= \frac{\Lambda_{\mathsf{f}}\mu_{\mathsf{f}} + \Lambda_s \tilde{z}_t}{\Lambda_{\mathsf{f}} + \Lambda_s},\\ \tilde{s}_t &= \bar{s}_t + (\Lambda_{\mathsf{f}} + \Lambda_s)^{-1/2}\xi_t,\\ \tilde{z}_{t+\Delta t} &= \tilde{s}_t + \Lambda_s^{-1/2}\epsilon_t,\end{aligned} \tag{S19}$$

where $\epsilon_t$ and $\xi_t$ are independent samples from a standard normal distribution.

In Eq. (S19), the mean and variance of $\tilde{s}_t$ and $\tilde{z}_t$ conditioned on $\mu_{\mathsf{f}}$ (the mean of the likelihood conveyed by feedforward input $\mathbf{u}^{\mathsf{f}}$, Eq. 19) in equilibrium ($t \to \infty$) are calculated as,

$$\begin{aligned}\langle \bar{s}|\mu_{\mathsf{f}}\rangle &= \mu_{\mathsf{f}}, \quad V(\bar{s}|\mu_{\mathsf{f}}) = \frac{\Lambda_s}{\Lambda_{\mathsf{f}}(\Lambda_{\mathsf{f}} + \Lambda_s)},\\ \langle \tilde{s}|\mu_{\mathsf{f}}\rangle &= \mu_{\mathsf{f}}, \quad V(\tilde{s}|\mu_{\mathsf{f}}) = \Lambda_{\mathsf{f}}^{-1},\\ \langle \tilde{z}|\mu_{\mathsf{f}}\rangle &= \mu_{\mathsf{f}}, \quad V(\tilde{z}|\mu_{\mathsf{f}}) = \Lambda_{\mathsf{f}}^{-1} + \Lambda_s^{-1}.\end{aligned} \tag{S20}$$

Here $\langle\cdot\rangle$ denotes an expectation over different realizations of $\epsilon_t$ and $\xi_t$.

##### 3.2 The generative model with latent stimuli organized in parallel

Now we analyze the equilibrium statistics of Gibbs sampling dynamics used to approximate the posterior of latent stimuli which are organized in parallel (Eq. 7). These sampling dynamics obey (Eqs. 8a and 8b in the main text):

$$p(\tilde{s}_1|\mathbf{u}_1^{\mathsf{f}}, \tilde{s}_{2,t-\Delta t}) \propto p(\mathbf{u}_1^{\mathsf{f}}|\tilde{s}_1)p(\tilde{s}_{2,t-\Delta t}|\tilde{s}_1), \tag{S21a}$$

$$\tilde{s}_{1t} \sim p(\tilde{s}_1|\mathbf{u}_1^{\mathsf{f}}, \tilde{s}_{2,t-\Delta t}). \tag{S21b}$$

Using the conditional distribution (Eq. S21a), the mean of the conditional distribution, $(\bar{s}_{1t}, \bar{s}_{2t})^\top$, must satisfy,

$$\begin{pmatrix}\Omega_1 & 0\\ 0 & \Omega_2\end{pmatrix}\begin{pmatrix}\bar{s}_{1t}\\ \bar{s}_{2t}\end{pmatrix} = \Lambda_s\begin{pmatrix}0 & 1\\ 1 & 0\end{pmatrix}\begin{pmatrix}\tilde{s}_{1,t-\Delta t}\\ \tilde{s}_{2,t-\Delta t}\end{pmatrix} + \begin{pmatrix}\Lambda_{\mathsf{f}1}\mu_{\mathsf{f}1}\\ \Lambda_{\mathsf{f}2}\mu_{\mathsf{f}2}\end{pmatrix},$$

where $\Omega_m = \Lambda_{\mathsf{f}m} + \Lambda_s$ $(m = 1, 2)$. Moreover, from the sampling step (Eq. S21b) we have

$$\begin{pmatrix}\tilde{s}_{1t}\\ \tilde{s}_{2t}\end{pmatrix} = \begin{pmatrix}\bar{s}_{1t}\\ \bar{s}_{2t}\end{pmatrix} + \begin{pmatrix}\Omega_1^{-1/2}\xi_{1t}\\ \Omega_2^{-1/2}\xi_{2t}\end{pmatrix}$$

Combining above two equations we derive the dynamics for $\bar{s}_{1t}$ and $\bar{s}_{2t}$.

$$\begin{pmatrix} \bar{s}_{1t} \\ \bar{s}_{2t} \end{pmatrix} = \Lambda_s \begin{pmatrix} 0 & \Omega_1^{-1} \\ \Omega_2^{-1} & 0 \end{pmatrix} \begin{pmatrix} \bar{s}_{1,t-\Delta t} \\ \bar{s}_{2,t-\Delta t} \end{pmatrix} + \Lambda_s \begin{pmatrix} \Omega_1^{-1}\Omega_2^{-1/2}\xi_{2t} \\ \Omega_2^{-1}\Omega_1^{-1/2}\xi_{1t} \end{pmatrix} + \begin{pmatrix} \Omega_1^{-1}\Lambda_{\mathsf{f}1}\mu_{\mathsf{f}1} \\ \Omega_2^{-1}\Lambda_{\mathsf{f}2}\mu_{\mathsf{f}2} \end{pmatrix} \tag{S22}$$

Then the mean of stimulus samples $\tilde{\mathbf{s}}_t = (\tilde{s}_{1t}, \tilde{s}_{2t})^\top$, and the mean of conditional distributions $\bar{\mathbf{s}}_t = (\bar{s}_{1t}, \bar{s}_{2t})^\top$ in equilibrium ($t \to \infty$) becomes

$$\langle \bar{\mathbf{s}}_t \rangle = \langle \tilde{\mathbf{s}}_t \rangle = \begin{pmatrix} \Omega_1 & -\Lambda_s \\ -\Lambda_s & \Omega_2 \end{pmatrix}^{-1} \begin{pmatrix} \Lambda_{\mathsf{f}1}\mu_{\mathsf{f}1} \\ \Lambda_{\mathsf{f}2}\mu_{\mathsf{f}2} \end{pmatrix}, \tag{S23}$$

where $\langle \cdot \rangle$ means the average over different realizations of $\xi_{1t}$ and $\xi_{2t}$. It can be checked that the mean of stimulus samples is the same as the posterior mean (Eq. 34).

The covariance of the mean generated from the conditional distribution is defined as, $\mathbf{\Sigma}(\bar{\mathbf{s}}_t) = \langle (\bar{\mathbf{s}}_t - \langle \bar{\mathbf{s}}_t \rangle)(\bar{\mathbf{s}}_t - \langle \bar{\mathbf{s}}_t \rangle)^\top \rangle$. Based on Eq. (S22), the discrete dynamics of $\mathbf{\Sigma}(\bar{s}_t)$ obey,

$$\mathbf{\Sigma}(\bar{s}_t) = \mathbf{A}\mathbf{\Sigma}(\bar{s}_{t-\Delta t})\mathbf{A}^\top + \mathbf{\Gamma} \tag{S24}$$

where

$$\mathbf{A} = \Lambda_s \begin{pmatrix} 0 & \Omega_1^{-1} \\ \Omega_2^{-1} & 0 \end{pmatrix}, \quad \mathbf{\Gamma} = \Lambda_s^2 \Omega_1^{-1}\Omega_2^{-1} \begin{pmatrix} \Omega_1^{-1} & 0 \\ 0 & \Omega_2^{-1} \end{pmatrix}.$$

In equilibrium, i.e. $\mathbf{\Sigma}(\bar{s}_t) = \mathbf{\Sigma}(\bar{s}_{t-\Delta t})$, we have that

$$\mathbf{\Sigma}(\bar{s}) \triangleq \mathbf{\Sigma}(\bar{s}_{t\to\infty}) = \frac{\Lambda_s^2}{\Omega_1\Omega_2 - \Lambda_s^2} \begin{pmatrix} \Omega_1^{-1} & 0 \\ 0 & \Omega_2^{-1} \end{pmatrix}. \tag{S25}$$

Furthermore, the covariance of stimulus samples in equilibrium $\mathbf{\Sigma}(\tilde{\mathbf{s}}) = \langle (\tilde{\mathbf{s}}_t - \langle \tilde{\mathbf{s}}_t \rangle)(\tilde{\mathbf{s}}_t - \langle \tilde{\mathbf{s}}_t \rangle)^\top \rangle|_{t\to\infty}$ becomes

$$\mathbf{\Sigma}(\tilde{\mathbf{s}}) = \mathbf{\Sigma}(\bar{\mathbf{s}}) + \begin{pmatrix} \Omega_1^{-1} & 0 \\ 0 & \Omega_2^{-1} \end{pmatrix} = \frac{1}{\Omega_1\Omega_2 - \Lambda_s^2} \begin{pmatrix} \Omega_2 & 0 \\ 0 & \Omega_1 \end{pmatrix}. \tag{S26}$$

It can be checked that the equilibrium variance of $\tilde{s}_{1t}$ and $\tilde{s}_{1t}$ are the same as the variance of the posterior distribution given in Eq. 34.

### 4 The neural responses distribution

#### 4.1 The single recurrent network

We present the derivation of the distribution of neuronal responses, $\mathbf{r}$, given an external stimulus feature $s$, i.e., $p(\mathbf{r}|s)$. For a fixed external stimulus, $s$, the neuronal response $\mathbf{r}$ fluctuates over time/trial due to both sensory transmission noise described by $p(\mathbf{u}^{\mathsf{f}}|s)$ (Eq. 23 and Fig. 2A, bottom), as well as the internally generated variability described by $p(\mathbf{r}|\mathbf{u}^{\mathsf{f}})$. Hence we have,

$$\begin{aligned} p(\mathbf{r}|s) &= \int p(\mathbf{r}|\mathbf{u}^{\mathsf{f}})p(\mathbf{u}^{\mathsf{f}}|s)d\mathbf{u}^{\mathsf{f}}, \\ &= \int \Big[ \int p(\mathbf{r}|\boldsymbol{\lambda})p(\boldsymbol{\lambda}|\mathbf{u}^{\mathsf{f}})d\boldsymbol{\lambda}\Big] p(\mathbf{u}^{\mathsf{f}}|s)d\mathbf{u}^{\mathsf{f}}. \end{aligned} \tag{S27}$$

Furthermore, internal variability, captured by $p(\mathbf{r}|\mathbf{u}^{\mathsf{f}})$, has two parts: One is $p(\mathbf{r}|\boldsymbol{\lambda})$ which describes the independent Poisson spike generation (Eq. 11); and the other is $p(\boldsymbol{\lambda}|\mathbf{u}^{\mathsf{f}})$ which describes the fluctuation of instantaneous firing rate, $\boldsymbol{\lambda}$, due to internal variability. Here $p(\boldsymbol{\lambda}|\mathbf{u}^{\mathsf{f}})$ is an unknown distribution in Eq. (S27), and we must compute it afterwards.

Since we are mainly interested in how stimulus is represented by neuronal activity, we only consider the covariability along the subspace defined by stimulus response, and ignore the covariability due to fluctuation along other directions. The instantaneous firing rate, $\boldsymbol{\lambda}_t$, can be approximated as a smooth Gaussian profile over $\bar{s}_t$, where $\bar{s}_t$ is the mean of the instantaneous conditional distribution (Eq. 12),

$$\boldsymbol{\lambda}_{tj}(\bar{s}_t) \approx R\exp[-(\bar{s}_t - \theta_j)^2/(2a^2)].$$

Assuming the variance of $\bar{s}$ (Eq. S20) is much smaller than the width of firing rate profile, i.e., $V(\tilde{s}|\mu_{\mathsf{f}}) \ll a$, the average of instantaneous firing rate $\boldsymbol{\lambda}_t$ over time can be approximated as a Gaussian profile whose mean is the average of $\bar{s}_t$ over time,

$$\begin{aligned} \langle \boldsymbol{\lambda}_t(\bar{s}_t)|\mathbf{u}^{\mathsf{f}}\rangle &\approx \boldsymbol{\lambda}(\langle \bar{s}_t|\mathbf{u}^{\mathsf{f}}\rangle), \\ &= R\exp[-(\mu_{\mathsf{f}} - \theta_j)^2/(2a^2)] \triangleq \mathbf{f}(\mu_{\mathsf{f}}). \end{aligned} \tag{S28}$$

The second equality in above equation comes from $\langle \bar{s}_t|\mathbf{u}^{\mathsf{f}}\rangle = \mu_{\mathsf{f}}$ as derived in Eq. (S20). Expanding the instantaneous firing rate $\boldsymbol{\lambda}_t(\bar{s})$ around the mean responses $\mathbf{f}(\mu_{\mathsf{f}})$,

$$\boldsymbol{\lambda}(\bar{s}_t) = \mathbf{f}(\mu_{\mathsf{f}}) + \mathbf{f}'_{\mu_{\mathsf{f}}}(\bar{s}_t - \mu_{\mathsf{f}}),$$

where $\mathbf{f}'_{\mu_{\mathsf{f}}} = d\mathbf{f}(\mu_{\mathsf{f}})/d\mu_{\mathsf{f}} = Ra^{-2}(\theta_j - \mu_{\mathsf{f}})\exp[-(\mu_{\mathsf{f}} - \theta_j)^2/(2a^2)]$. The covariance of the instantaneous firing rate becomes

$$\begin{aligned}\Sigma[\boldsymbol{\lambda}(\bar{s})] &= \left\langle [\boldsymbol{\lambda}(\bar{s}_t) - \mathbf{f}(\mu_\mathsf{f})][\boldsymbol{\lambda}(\bar{s}_t) - \mathbf{f}(\mu_\mathsf{f})]^\top \right\rangle, \\ &= \left\langle (\bar{s}_t - \mu_\mathsf{f})^2 \right\rangle \mathbf{f}'_{\mu_\mathsf{f}} \mathbf{f}'^\top_{\mu_\mathsf{f}}, \\ &= V(\bar{s}|\mu_\mathsf{f}) \mathbf{f}'_{\mu_\mathsf{f}} \mathbf{f}'^\top_{\mu_\mathsf{f}}. \end{aligned} \tag{S29}$$

$V(\bar{s}|\mu_\mathsf{f})$ is the variance of $\bar{s}$ given the feedforward input $\mathbf{u}^\mathsf{f}$, as derived in Eq. (S20). The covariance structure $\mathbf{f}'_{\mu_\mathsf{f}} \mathbf{f}'^\top_{\mu_\mathsf{f}}$ captures the covariability due to firing rate fluctuations along the stimulus subspace, which is often termed differential (noise) correlations [3, 4]. With the Gaussian tuning of $\mathbf{f}(\mu_\mathsf{f})$ (Eq. S28), $\mathbf{f}'_{\mu_\mathsf{f}} \mathbf{f}'^\top_{\mu_\mathsf{f}}$ exhibits anti-symmetric structure over $\mu_\mathsf{f}$, i.e.,

$$\left[\mathbf{f}'_{\mu_\mathsf{f}} \mathbf{f}'^\top_{\mu_\mathsf{f}}\right]_{ij} \propto (\theta_i - \mu_\mathsf{f})(\theta_j - \mu_\mathsf{f}) \exp[-(\mu_\mathsf{f} - \theta_i)^2/(2a^2)] \exp[-(\mu_\mathsf{f} - \theta_j)^2/(2a^2)],$$

(see Fig. 8B). Combining Eqs. (S28 and S29) together, $p(\boldsymbol{\lambda}|\mathbf{u}^\mathsf{f})$ can be approximated as a multivariate normal distribution,

$$p(\boldsymbol{\lambda}|\mathbf{u}^\mathsf{f}) \approx \mathcal{N}\left[\boldsymbol{\lambda} \middle| \mathbf{f}(\mu_\mathsf{f}), V(\bar{s}|\mu_\mathsf{f}) \mathbf{f}'_{\mu_\mathsf{f}} \mathbf{f}'^\top_{\mu_\mathsf{f}}\right], \tag{S30}$$

where,

$$\mathbf{f}_j(\mu_\mathsf{f}) = R \exp[-(\mu_\mathsf{f} - \theta_j)^2/2a^2], \tag{S31}$$

$$V(\bar{s}|\mu_\mathsf{f}) = \frac{\Lambda_s}{\Lambda_\mathsf{f}(\Lambda_\mathsf{f} + \Lambda_s)} = a^2 n_\mathsf{f}^{-1} w_E^*. \tag{S32}$$

Here we have derived the Eq. (44) shown in the main text. The 2nd equality in Eq. (S32) is obtained by using Eq. (6) in the main text. Furthermore, by approximating the independent Poisson distributions $p(\mathbf{r}|\boldsymbol{\lambda})$ and $p(\mathbf{u}^\mathsf{f}|s)$ as multivariate normal distribution, i.e.,

$$p(\mathbf{r}|\boldsymbol{\lambda}) \approx \mathcal{N}[\mathbf{r}|\boldsymbol{\lambda}(\bar{s}), \mathrm{diag}(\boldsymbol{\lambda}(\bar{s}))],$$

the distribution $p(\boldsymbol{r}|\mathbf{u}^\mathsf{f})$ can be computed as

$$\begin{aligned} p(\mathbf{r}|\mathbf{u}^\mathsf{f}) &= \int p(\mathbf{r}|\boldsymbol{\lambda}) p(\boldsymbol{\lambda}|\mathbf{u}^\mathsf{f}) d\boldsymbol{\lambda}, \\ &= \mathcal{N}\left[\mathbf{r} \middle| \mathbf{f}(\mu_\mathsf{f}), \mathrm{diag}(\boldsymbol{\lambda}(\bar{s})) + V(\bar{s}|\mu_\mathsf{f}) \mathbf{f}'_{\mu_\mathsf{f}} \mathbf{f}'^\top_{\mu_\mathsf{f}}\right]. \end{aligned} \tag{S33}$$

Compared with $p(\boldsymbol{\lambda}|\mathbf{u}^\mathsf{f})$ (Eq. S30), we see the covariance of $p(\mathbf{r}|\mathbf{u}^\mathsf{f})$ (Eq. S33) has an extra term, i.e., $\mathrm{diag}(\boldsymbol{\lambda}(\bar{s}))$, which is a diagonal matrix denoting the independent Poisson spiking variability. Finally, substituting Eqs. (S30) and (23) into Eq. (S27), shows that the response distribution conditioned on the external stimulus feature $s$, $p(\mathbf{r}|s)$, has the form

$$p(\mathbf{r}|s) \approx \mathcal{N}\big[\mathbf{r}\big|\mathbf{f}(s), \mathrm{diag}(\mathbf{f}(s)) + V(\bar{s}|s)\mathbf{f}'_s\mathbf{f}_s'^{\top}\big]. \tag{S34}$$

Here the variance $V(\bar{s}|s)$ in the stimulus subspace is a mixture of internal variability due to Poisson spike discharge and sensory noise. It follows from Eqs. S20 and 23 that

$$\begin{aligned} V(\bar{s}|s) &= V(\bar{s}|\mu_{\mathsf{f}}) + V(\mu_{\mathsf{f}}|s), \\ &= \frac{\Lambda_s}{\Lambda_{\mathsf{f}}(\Lambda_{\mathsf{f}} + \Lambda_s)} + \frac{1}{\Lambda_{\mathsf{f}}}, \\ &= a^2 n_{\mathsf{f}}^{-1}(w_E^* + 1). \end{aligned} \tag{S35}$$

The second term, $V(\mu_{\mathsf{f}}|s)$, represents fluctuations in neuronal activity in the stimulus subspace due to the sensory transmission noise (Eq. 23).

#### 4.2 The coupled networks

We extend the derivation shown from Eq. (S27) to Eq. (S35) to calculate the response distribution for the coupled network used to infer latent variables organized in parallel (Fig. 6A). We only present the results for network 1 for example, as those for network 2 can be obtained by changing indices Following the analysis above we consider only neuronal fluctuations along the stimulus subspace,

$$p(\mathbf{r}_1|\mathbf{s}) \approx \mathcal{N}[\mathbf{r}_1|\mathbf{f}(\langle\bar{s}_1\rangle), V(\bar{s}_1)\mathbf{f}'_s\mathbf{f}_s'^{\top}]. \tag{S36}$$

Here $\mathbf{s} = (s_1, s_2)^\top$ are the external stimuli presented to the network (Eq. 7), $\langle\bar{s}_1\rangle$ and $V(\bar{s}_1)$ are the mean and variance of $\bar{s}_{1t}$ in equilibrium, and $\bar{s}_{1t}$ denotes the mean of conditional distribution $p(\tilde{s}_1|\mathbf{u}_1^{\mathsf{f}}, \tilde{s}_{2,t-\Delta t})$ (Eq. 8a).

Based on the results shown in Eqs. (S23 and S25), the mean and variance of $\mathbf{s}_m$ given external stimuli $\mathbf{s}$ are calculated as,

$$\begin{aligned} \langle\bar{s}_1|\mathbf{s}\rangle &= \frac{(\Lambda_2^{-1} + \Lambda_s^{-1})s_1 + \Lambda_1^{-1}s_2}{\Lambda_1^{-1} + \Lambda_2^{-1} + \Lambda_s^{-1}}, \\ V(\bar{s}_1|\mathbf{s}) &= \frac{\Lambda_1^{-1}\Lambda_2^{-1}}{\Lambda_1^{-1} + \Lambda_2^{-1} + \Lambda_s^{-1}} \frac{\Lambda_1^{-1}}{\Lambda_1^{-1} + \Lambda_s^{-1}} + \Lambda_1^{-1}. \end{aligned} \tag{S37}$$

The variance $V(\bar{s}_1|\mathbf{s})$ characterizes the total amount of differential correlations in the neuronal response to a fixed stimuli $\mathbf{s}$. We see that differential correlations decrease with the precision $\Lambda_1$ which is determined by the feedforward input rate (Eq. 20). Moreover, differential correlations increase with the precision $\Lambda_s$, which determines the prior correlation between two stimuli (Eq. 7) and is stored by the coupling weight between the two networks (Eq. 40). These results are confirmed by numerical simulations (Fig. S5).

#### 5 Invertible linear transformations do not degrade linear Fisher information

A network that linearly transforms its inputs does not degrade linear Fisher information. On the other hand, internally generated differential correlations will decrease linear Fisher information. To prove this, we write the activity of cells in a linear network in response to a stimulus $s$ as,

$$\mathbf{r}(s) = \mathbf{A}\mathbf{u}^{\mathsf{f}}(s) + \epsilon\boldsymbol{\eta}.$$

Here $\mathbf{A}$ is the linear transformation matrix and is determined by the recurrent connections in the network, $\mathbf{u}^{\mathsf{f}}(s)$ is the feedforward input defined in Eq. (18). The term $\epsilon\boldsymbol{\eta}$, captures the internally generated differential correlations with amplitude controlled by $\epsilon$ (Eq. 9). The random variable $\boldsymbol{\eta}$ has zero mean, and covariance equal to $\Sigma_{\boldsymbol{\eta}} = \mathbf{f}'_s\mathbf{f}'^{\top}_s$, with $\mathbf{f}'_s = d\mathbf{f}(s)/ds$, and $\mathbf{f}(s) = \langle \mathbf{r}(s)\rangle$ is the tuning of network responses (Eq. 9). When the matrix $\mathbf{A}$ is *invertible*, the linear Fisher information of $s$ in neuronal response $\mathbf{r}$ is,

$$\begin{aligned}
\mathcal{I}_{\mathbf{r}}(s) &= \mathbf{f}'^{\top}_s \Sigma_{\mathbf{r}}^{-1}\mathbf{f}'_s, \\
&= \left(\mathbf{A}\langle \mathbf{u}^{\mathsf{f}}\rangle'_s\right)^{\top}\left[\mathbf{A}\Sigma_{\mathbf{u}^{\mathsf{f}}}\mathbf{A}^{\top} + \epsilon\mathbf{f}'_s\mathbf{f}'^{\top}_s\right]^{-1}\mathbf{A}\langle \mathbf{u}^{\mathsf{f}}\rangle'_s, \\
&= \langle \mathbf{u}^{\mathsf{f}}\rangle'^{\top}_s\left[\Sigma_{\mathbf{u}^{\mathsf{f}}} + \epsilon\langle \mathbf{u}^{\mathsf{f}}\rangle'_s\langle \mathbf{u}^{\mathsf{f}}\rangle'^{\top}_s\right]^{-1}\langle \mathbf{u}^{\mathsf{f}}\rangle'_s,
\end{aligned}$$

where $\langle \mathbf{u}^{\mathsf{f}}\rangle$ is the tuning of feedforward inputs, $\mathbf{u}^{\mathsf{f}}$ (Eq. 18).

Thus the linear Fisher information does not depend on $\mathbf{A}$, and hence a linear invertible transformation does not decrease the linear Fisher information. By using the matrix inverse lemma we find that

$$\left[\Sigma_{\mathbf{u}^{\mathsf{f}}} + \epsilon\langle \mathbf{u}^{\mathsf{f}}\rangle'_s\langle \mathbf{u}^{\mathsf{f}}\rangle'^{\top}_s\right]^{-1} = \Sigma_{\mathbf{u}^{\mathsf{f}}}^{-1} - \frac{\epsilon}{1+\epsilon\langle \mathbf{u}^{\mathsf{f}}\rangle'^{\top}_s\Sigma_{\mathbf{u}^{\mathsf{f}}}^{-1}\langle \mathbf{u}^{\mathsf{f}}\rangle'_s}\Sigma_{\mathbf{u}^{\mathsf{f}}}^{-1}\langle \mathbf{u}^{\mathsf{f}}\rangle'_s\langle \mathbf{u}^{\mathsf{f}}\rangle'^{\top}_s\Sigma_{\mathbf{u}^{\mathsf{f}}}^{-1},$$

the linear Fisher information of stimulus $s$ in $\mathbf{r}$ can be computed as,

$$\mathcal{I}_{\mathbf{r}}(s) = \frac{\langle \mathbf{u}^{\mathsf{f}}\rangle'^{\top}_s\Sigma_{\mathbf{u}^{\mathsf{f}}}^{-1}\langle \mathbf{u}^{\mathsf{f}}\rangle'_s,}{1+\epsilon\langle \mathbf{u}^{\mathsf{f}}\rangle'^{\top}_s\Sigma_{\mathbf{u}^{\mathsf{f}}}^{-1}\langle \mathbf{u}^{\mathsf{f}}\rangle'_s} = \frac{\mathcal{I}_{\mathbf{u}^{\mathsf{f}}}(s)}{1+\epsilon\mathcal{I}_{\mathbf{u}^{\mathsf{f}}}(s)}. \tag{S38}$$

If the linear network doesn't internally generate differential correlation, i.e., $\epsilon = 0$, the linear Fisher information in the network response is the same as the the information inherited from feedforward input, i.e., $\mathcal{I}_{\mathbf{r}}(s) = \mathcal{I}_{\mathbf{u}^{\mathsf{f}}}(s)$. Hence internal differential correlations degrade linear Fisher information,

| Symbol | Description | Typical values (range) |
|---|---|---|
| | **Generative model** | |
| $z$ | Context (scalar variable) | $(-180°, 180°]$ |
| $s_m$ | Stimulus $m$ (scalar variable) | $(-180°, 180°]$ |
| $\mathbf{u}^\mathsf{f}$ | Feedforward spiking inputs (vector) | |
| $U^\mathsf{f}$ | Peak feedforward input rate | $[0, 50]$Hz |
| $\Lambda_s$ | The prior precision | |
| $\Lambda_\mathsf{f}$ | The likelihood precision | |
| | **Network model** | |
| $N_E$ | Number of excitatory neurons | 180 |
| $N_I$ | Number of inhibitory neurons | 45 |
| $a$ | Tuning width | 40° |
| $\tau_d$ | Synaptic decaying time constant | 2 ms |
| $\mathbf{r}_t$ | Neuronal spikes at time $t$ (vector) | |
| $\boldsymbol{\lambda}_t$ | Firing rate at time $t$ (vector) | |
| $\mathbf{u}_t^\mathsf{r}$ | Recurrent input at time $t$ (vector) | |
| $U^\mathsf{r}$ | Peak value of recurrent input | |
| $n_\mathsf{f}$ | The total spike count of $\mathbf{u}^\mathsf{f}$ to all neurons (scalar variable) | |
| $n_\mathsf{r}$ | The sum of recurrent input (scalar variable) | |
| $n_{\boldsymbol{\lambda}}$ | The sum of population firing rate (scalar variable) | |
| $w_E$ | The peak recurrent weight between excitatory neurons | $[0, 5 \times 10^{-3}]$ |
| $w_I$ | The inhibitory synaptic weight | $5w_E$ |
| $w_{I\mathsf{f}}$ | The feedforward weight to inhibitory neurons | 0.8 |
| $w_{EE}^{mn}$ | The E weight across networks in coupled circuit | 0.01 |
| $\theta_j$ | The preferred stimulus of $j^{\text{th}}$ excitatory neuron | |
| $dt$ | Time step in numerical simulation | 0.1 ms |
| $T_{decode}$ | The time window of decoding a stimulus sample | 20 ms |

Table 1: Notations and parameters in the model

but any linear invertible transformations by the network do not.

### 6 Simulation details and model parameters

#### 6.1 Parameters used in network simulation

Table 1 lists the notations and typical parameters used in the model and network simulations. The results shown in the main text Figures are obtained with the typical parameters if not mentioned otherwise.

Below summarizes some particular parameters used in figures.

- Figure 4: $w_E = 0.02$
- Figure 5: $U^\mathsf{f} = 50$Hz.
- Figure 6B-D: $w_{EE}^{mn} = 0.05$.

- Figure 7D: $U^{\mathsf{f}1} = U^{\mathsf{f}2} = 30$Hz
- Figure 7E: $w_{EE}^{mn} = 0.01$, $U^{\mathsf{f}2} = 25$Hz.
- Figure 7F and G: $U_m^{\mathsf{f}} \sim \mathcal{U}[0, 50\text{Hz}]$, $\mu_{\mathsf{f}m} \sim \mathcal{U}[-10^\circ, 10^\circ]$, $w_{EE}^{mn} \sim \mathcal{U}[0, 0.05]$. Simulated under 1000 randomly generated parameters.
- Figure 8C and 8E: $U^{\mathsf{f}} = 30$Hz.
- Figure 8D: $w_E = 1 \times 10^{-3}$.

#### 6.2 The single recurrent network

##### 6.2.1 Estimating the sampling distribution and mutual information

To estimate the sampling distribution and the mutual information in the network responses, we simulated the network with a fixed feedforward input $\mathbf{u}^{\mathsf{f}}$ but which is randomly generated given the stimulus $s$ in the external world. Without loss of generality we set the external stimulus $s$ to zero, since the feedforward inputs and network responses are translation-invariant.

The spiking network (Eqs. 47-50) was simulated for 102 seconds, and the neuronal responses during the first 2 seconds were discarded in all further analysis. In each time window lasting 20 ms, the samples of the stimulus and context were read out by population vector from the spiking activities of E neurons, $\mathbf{r}^E$, and recurrent input of E neurons lasting over the time window, $\mathbf{u}^{E\mathsf{r}}$ respectively,

$$\tilde{s}_t = \textstyle\sum_j \mathbf{r}_j^E(t)\theta_j / \sum_j \mathbf{r}_j^E(t), \qquad \tilde{z}_t = \sum_j \mathbf{u}_j^{E\mathsf{r}}(t)\theta_j / \sum_j \mathbf{u}_j^{E\mathsf{r}}(t).$$

The duration of the decoding time window is not critical in obtaining the sampling distribution; it only needs to be long enough to allow a neuron in the population fires a spike in this time window. The joint distribution of samples $\tilde{s}$ and $\tilde{z}$ is approximated as a bivariate normal distribution, and its mean, $\boldsymbol{\mu}_q$, and precision matrix, $\mathbf{K}_q$, are computed.

Estimating the mutual information (Eq. 43) between latent variables, i.e., $s$ and $z$, and the feedforward input $\mathbf{u}^{\mathsf{f}}$ requires the posterior distribution (Eq. 24). Computing the posterior (Eq. 24) requires the likelihood function and prior distribution. The mean, $\mu_{\mathsf{f}}$, and precision, $\Lambda_{\mathsf{f}}$, of the likelihood were directly read out from the feedforward input (Eq. 20). Meanwhile, the prior precision, $\Lambda_s$, was pre-assigned. Substituting the parameters of the sampling distribution, i.e., $\boldsymbol{\mu}_q$ and $\mathbf{K}_q$, and the parameters of posterior, i.e., $\boldsymbol{\mu}_p$ and $\mathbf{K}_p$, into Eq. (43), permits the mutual information to be calculated.

###### 6.2.2 Estimating the linear Fisher information

We estimated the linear Fisher information of the stimulus $s$ from the responses of E neurons, $\mathbf{r}^E$, using a bias-correction algorithm [5]. Given a pair of external stimuli $s_\pm = \pm 1^\circ$, we randomly generate the feedforward inputs in every time step (Eq. 18) and simulate the network for $T = 1\times 10^4$ trials with each trial lasting 200ms. The mean and covariance of E neurons' responses $\mathbf{r}^E$ under each stimulus are recorded. Then the bias-correction estimate of linear Fisher information is,

$$\mathcal{I}_{bc}(s) = \frac{d\langle \mathbf{r}^E\rangle}{ds}^\top \Sigma(\mathbf{r}^E)^{-1} \frac{d\langle \mathbf{r}^E\rangle}{ds} \frac{2T-N-3}{2T-2} - \frac{2N}{Tds^2}, \tag{S39}$$

where $d\langle \mathbf{r}\rangle = \langle \mathbf{r}^E_+\rangle - \langle \mathbf{r}^E_-\rangle$, $ds = s_+ - s_-$, and $\langle\cdot\rangle$ represents the average over trials. $\Sigma(\mathbf{r}^E)$ is the covariance matrix of E neurons' responses. $N$ is the number of neurons in the network.

###### 6.2.3 Comparing the sampling precision with posterior precision

Due to symmetry in the recurrent coupling in the neural population (Fig. 4A), the means of stimulus samples and context samples are always the same, and are the same as the posterior mean (Eq. 24). Therefore, we only compare the sampling precision with the posterior precision in order to evaluate the accuracy of network sampling.

To obtain the posterior precision matrix, we first read out the likelihood directly from the feedforward inputs (Eq. 20). Since we do not know the value of the prior precision $\Lambda_s$ stored in the network, given each sampling distribution generated by the network, we numerically obtained the value of $\Lambda_s$ which minimizes the KL divergence from the posterior to the sampling distribution. Given this $\Lambda_s$, the prediction of posterior precision is computed as $\mathbf{K}_p = \mathbf{\Lambda}_s + \mathbf{\Lambda}_\mathsf{f}$ (Eq. 24) which is then compared with actual sampling precision (Fig. S4).

It is worth noting that the searched prior precision $\Lambda_s$ is a *subjective* prior, which reflects the prior stored in the recurrent network and may change with input.

##### 6.3 The coupled networks

In order to estimate the sampling distribution and the mutual information in the coupled networks, we simulate the network model for 102 seconds in response to fixed feedforward spiking inputs which are randomly generated from the generative model (Eq. 33). Then the network responses in first 2 seconds were discarded for further analysis. On each decoding time window of 20ms, the samples of each stimulus is read out by population vector from E neurons in corresponding network,

$$\tilde{s}_{mt} = \textstyle\sum_j \mathbf{r}^E_{mj}(t)\theta_j / \sum_j \mathbf{r}^E_{mj}(t), \quad (m=1,2). \tag{S40}$$

For these simulations we compute the mean, $\boldsymbol{\mu}_q$, and covariance matrix, $\mathbf{K}_q$, of stimulus samples, which are then be substituted into Eq. (43) to compute the mutual information.

### 7 Appendix

#### 7.1 Computing the likelihood using Laplace's method

We present the derivation used to compute the $\hat{n}_{\mathbf{r}}$ and H defined in Eq. (S7) in order to approximate the integral in Eq. (S6) by using Laplace's method [1]. To simplify notations, we denote $\mathcal{L} \equiv p(\tilde{s}, n_{\mathbf{r}} | \bar{s}, n_{\boldsymbol{\lambda}})$, and then,

$$\ln \mathcal{L} \;=\; \frac{1}{2}\ln n_{\mathbf{r}} - \frac{n_{\mathbf{r}}}{2a^2}(\tilde{s}-\bar{s})^2 - \frac{(n_{\mathbf{r}} - n_{\boldsymbol{\lambda}})^2}{2n_{\boldsymbol{\lambda}}}. \tag{A1}$$

$\hat{n}_{\mathbf{r}}$ can be found by taking the derivative of $\mathcal{L}$ over $n_{\mathbf{r}}$ to be zero,

$$\frac{\partial \ln \mathcal{L}}{\partial n_{\mathbf{r}}} \;=\; \frac{1}{2n_{\mathbf{r}}} - \frac{(\tilde{s}-\bar{s})^2}{2a^2} - \frac{n_{\mathbf{r}} - n_{\boldsymbol{\lambda}}}{n_{\boldsymbol{\lambda}}} = 0,$$

then the $\hat{n}_{\mathbf{r}}$ is computed as

$$\begin{aligned} \hat{n}_{\mathbf{r}} \;&=\; \left[1 - \frac{(\tilde{s}-\bar{s})^2}{2a^2}\right]\frac{n_{\boldsymbol{\lambda}}}{2} + \frac{1}{2}\sqrt{\left[1 - \frac{(\tilde{s}-\bar{s})^2}{2a^2}\right]^2 n_{\boldsymbol{\lambda}}^2 + 2n_{\boldsymbol{\lambda}}}, \\ &\approx\; \left[1 - \frac{(\tilde{s}-\bar{s})^2}{2a^2}\right] n_{\boldsymbol{\lambda}} + \mathcal{O}\big(n_{\boldsymbol{\lambda}}^{1/2}\big). \end{aligned} \tag{A2}$$

To gain insight, we simplify the above equation by omitting the term $2n_{\boldsymbol{\lambda}}$ inside the square root function (second line of Eq. A2). This approximation works well when $n_{\boldsymbol{\lambda}}$ is large enough. The negative Hessian matrix is,

$$\begin{aligned} \frac{\partial^2 \mathcal{L}}{\partial n_{\mathbf{r}}^2}\Big|_{n_{\mathbf{r}}=\hat{n}_{\mathbf{r}}} \;&=\; -\frac{1}{2\hat{n}_{\mathbf{r}}^2} - \frac{1}{n_{\boldsymbol{\lambda}}}, \\ &=\; -\frac{1}{2\big[1 - \frac{(\tilde{s}-\bar{s})^2}{2a^2}\big]^2 n_{\boldsymbol{\lambda}}^2} - \frac{1}{n_{\boldsymbol{\lambda}}}, \\ &\approx\; -\frac{1}{n_{\boldsymbol{\lambda}}} + \mathcal{O}\big(n_{\boldsymbol{\lambda}}^{-2}\big). \end{aligned} \tag{A3}$$

This approximation is also considered under the large $n_{\boldsymbol{\lambda}}$ limit where the omitted term is a order smaller than $1/n_{\boldsymbol{\lambda}}$.

Substituting Eqs. (A2 and A3) back into Eq. (S6), we get an unnormalized distribution of stimulus sample $\tilde{s}$,

$$p(\tilde{s}|\bar{s}, n_{\boldsymbol{\lambda}}) \propto \sqrt{\hat{n}_{\mathbf{r}}(\tilde{s})}\,\exp\Big[-\frac{\hat{n}_{\mathbf{r}}(\tilde{s})}{2a^2}(\tilde{s}-\bar{s})^2\Big]\exp\Big[-\frac{(\hat{n}_{\mathbf{r}}(\tilde{s}) - n_{\boldsymbol{\lambda}})^2}{2n_{\boldsymbol{\lambda}}}\Big]. \tag{A4}$$

Since this distribution is complicated, we again approximate it by a Gaussian distribution. It is easy to see $p(\tilde{s}|\bar{s}, n_{\boldsymbol{\lambda}})$ is a symmetric distribution over its center $\tilde{s} = \bar{s}$, and it is a mixture of

Gaussian distributions with different width (Fig. S3A). Next, we compute the Hessian of $p(\tilde{s}|\bar{s}, n_{\boldsymbol{\lambda}})$ at it peak location, i.e., $\tilde{s} = \bar{s}$, which is the precision of the approximated Gaussian distribution of $p(\tilde{s}|\bar{s}, n_{\boldsymbol{\lambda}})$. Denote by $\mathcal{L}(\tilde{s}) = p(\tilde{s}|\bar{s}, n_{\boldsymbol{\lambda}})$ to simplify notations, and we have,

$$\ln \mathcal{L}(\tilde{s}) = \frac{1}{2} \ln \hat{n}_{\mathbf{r}}(\tilde{s}) - \frac{\hat{n}_{\mathbf{r}}(\tilde{s})}{2a^2}(\tilde{s} - \bar{s})^2 - \frac{(\hat{n}_{\mathbf{r}}(\tilde{s}) - n_{\boldsymbol{\lambda}})^2}{2n_{\boldsymbol{\lambda}}}. \tag{A5}$$

The Hessian of $\mathcal{L}(\tilde{s})$ is calculated as

$$\begin{aligned}
\frac{\partial \ln \mathcal{L}(\tilde{s})}{\partial \tilde{s}} &= \frac{1}{2\hat{n}_{\mathbf{r}}(\tilde{s})} \frac{\partial \hat{n}_{\mathbf{r}}(\tilde{s})}{\partial \tilde{s}} - \frac{(\tilde{s} - \bar{s})^2}{2a^2} \frac{\partial \hat{n}_{\mathbf{r}}(\tilde{s})}{\partial \tilde{s}} - \frac{\hat{n}_{\mathbf{r}}(\tilde{s})}{a^2}(\tilde{s} - \bar{s}) - \frac{\hat{n}_{\mathbf{r}}(\tilde{s}) - n_{\boldsymbol{\lambda}}}{n_{\boldsymbol{\lambda}}} \frac{\partial \hat{n}_{\mathbf{r}}(\tilde{s})}{\partial \tilde{s}}, \\
\frac{\partial^2 \ln \mathcal{L}(\tilde{s})}{\partial \tilde{s}^2} &= -\left( \frac{1}{2\hat{n}_{\mathbf{r}}(\tilde{s})^2} + \frac{1}{n_{\boldsymbol{\lambda}}} \right) \left( \frac{\partial \hat{n}_{\mathbf{r}}(\tilde{s})}{\partial \tilde{s}} \right)^2 + \left[ \frac{1}{2\hat{n}_{\mathbf{r}}(\tilde{s})} - \frac{(\tilde{s} - \bar{s})^2}{2a^2} - \frac{\hat{n}_{\mathbf{r}}(\tilde{s})}{n_{\boldsymbol{\lambda}}} + 1 \right] \frac{\partial^2 \hat{n}_{\mathbf{r}}(\tilde{s})}{\partial \tilde{s}^2} \\
&\quad - 2\frac{(\tilde{s} - \bar{s})}{a^2} \frac{\partial \hat{n}_{\mathbf{r}}(\tilde{s})}{\partial \tilde{s}} - \frac{1}{a^2} \hat{n}_{\mathbf{r}}(\tilde{s}).
\end{aligned} \tag{A6}$$

Finally, the derivative of $\hat{n}_{\mathbf{r}}(\tilde{s})$ over $\tilde{s}$ is

$$\frac{\partial \hat{n}_{\mathbf{r}}(\tilde{s})}{\partial \tilde{s}} = \frac{n_{\boldsymbol{\lambda}}}{a^2}(\bar{s} - \tilde{s}), \quad \frac{\partial^2 \hat{n}(\tilde{s})}{\partial \tilde{s}^2} = -\frac{n_{\boldsymbol{\lambda}}}{a^2}. \tag{A7}$$

Substituting Eq. (A7) into Eq. (A6), the the Hessian of $\mathcal{L}(\tilde{s})$ at $\tilde{s} = \bar{s}$ can be calculated as,

$$\left. \frac{\partial^2 \ln \mathcal{L}(\tilde{s})}{\partial \tilde{s}^2} \right|_{\tilde{s} = \bar{s}} = -a^{-2} \left( \frac{1}{2} + n_{\boldsymbol{\lambda}} \right) \approx -a^{-2} n_{\boldsymbol{\lambda}}. \tag{A8}$$

We ignore the $1/2$ in the above equation since it is much smaller than $n_{\boldsymbol{\lambda}}$ when $n_{\boldsymbol{\lambda}}$ is large. Finally, the likelihood can be approximated as a Gaussian distribution

$$p(\tilde{s}|\bar{s}, n_{\boldsymbol{\lambda}}) \approx \mathcal{N}\left(\tilde{s}|\bar{s}, a^2 n_{\boldsymbol{\lambda}}^{-1}\right), \tag{A9}$$

which turns to be Eq. (S8).
